# A dynamic view of histone tails interaction with clustered abasic sites in a nucleosome core particle

**DOI:** 10.1101/2021.02.16.431417

**Authors:** Emmanuelle Bignon, Natacha Gillet, Tao Jiang, Christophe Morell, Elise Dumont

**Affiliations:** Univ Lyon, ENS de Lyon, CNRS UMR 5182, Laboratoire de Chimie, F69342, Lyon, France; Université de Lyon, Institut des Sciences Analytiques, UMR 5280 CNRS, Université Claude Bernard Lyon 1, 5 rue de la Doua, 69100 Villeurbanne, France; Institut Universitaire de France, 5 rue Descartes, 75005 Paris, France

## Abstract

Apurinic/apyrimidinic sites are the most common DNA damage under physiological conditions. Yet, their structural and dynamical behavior within nucleosome core particles has just begun to be investigated, and show dramatic differences with the one of abasic sites in B-DNA. Clusters of two or more abasic sites are repaired even less efficiently and hence constitute hotspots of high mutagenicity notably due to enhanced double-strand breaks formation. Based on a X-ray structure of a 146-bp DNA wrapped onto a histone core, we investigate the structural behavior of two bistranded abasic sites positioned at mutational hotspots along microsecond-range molecular dynamics simulations. Our simulations allow us to probe histone tails interactions at clustered abasic sites locations, with a definitive assignment of the key residues involved in the NCP-catalyzed formation of DNA–protein cross-linking in line with recent experimental findings, and pave the way towards a systematic assessment of histone tails response to DNA lesions.

---

Apurinic/apyrimidinic (Ap) sites are ubiquitous DNA damages, ^1^ with 10,000 such lesions produced per cell per day. Ap sites are reactive species, which can lead to more complex lesions and strand scission upon condensation with proximal amino groups if not correctly repaired. The formation and outcome of abasic sites within B-DNA has been intensively studied to delineate their processing by endonucleases (human APE1 and bacterial Nfo ^2^). Upon exposure to *γ* radiations and other damaging agents, accumulation of proximal abasic sites can give rise to multiple damaged sites within one or two helical turns of DNA. ^3,4^ Such clustered lesions can lead to the formation of highly deleterious double strand breaks (DSBs) via DNA-protein crosslinks (DPCs) formation with basic amino acids. ^5^ Whereas a large part of the chemistry of Ap sites has been investigated in naked oligonucleotides, their actual environment corresponds to a larger nucleosomal DNA fragment (146-bp N-DNA) wrapped onto an histone core that consists of a dimer of proteic tetramers – H2A, H2B, H3, and H4, shown in Figure 1.

**Figure 1.**
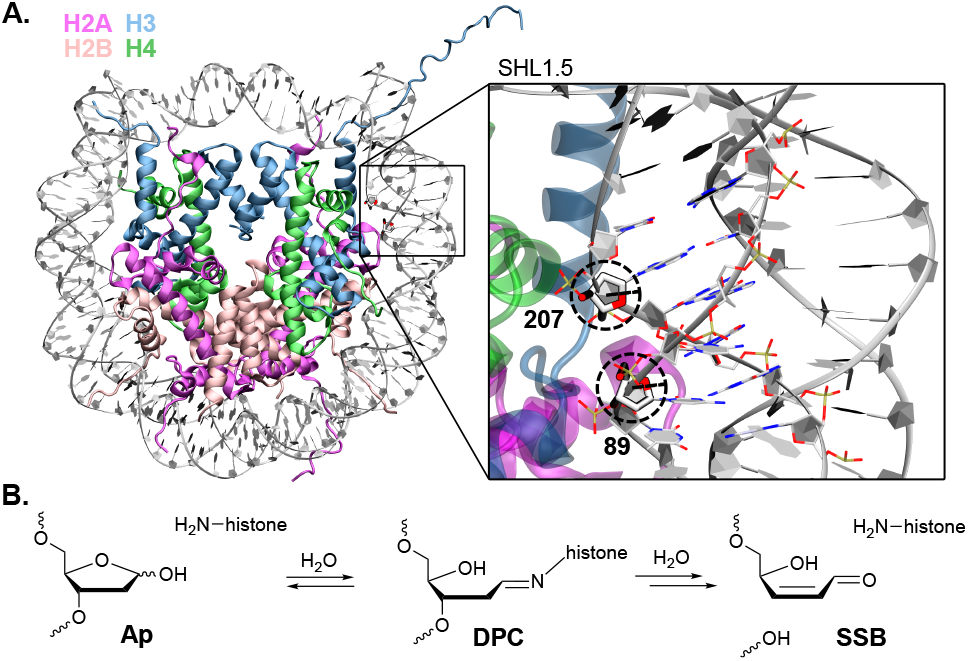
Location and reactivity of the clustered Ap sites at positions 89 and 207 within the nucleosome core particle (PDB ID 1AOI). (**A**) Positions 89 and 207 that are mutated *in silico* to Ap sites are located at super-helical location 1.5. (**B**) The Ap site reaction with a histone basic residue result in a DNA-protein cross-linking (DPC) via the formation of a Schiff base, and can ultimately be hydrolyzed into a single strand break (SSB).

As embedded in a nucleosome core particle (NCP), the helix experiences strong electrostatic interactions with proximal positively-charged residues and a mechanical constraint due to the coiling/wrapping that play a role in redefining Ap chemistry. Whereas the repair of Ap sites and other lesions can be mostly accounted for by a well-celebrated rotation effect, ^6,7^ the processing of NCP-catalyzed DPC and strand breaks from Ap sites is more complex. The NCP environment affects the lifetime and outcome of Ap sites, with various possibilities to undergo Schiff base formation in the “catalytic” environment provided by the histone core, ^8^ ultimately leading to strand scission resulting from *β*-elimination – see Figure 1-B. Kinetics of strand breaks are much higher at abasic sites embedded in a NCPs than in duplexes, with reactivity of Ap sites raised by up to 5-fold in NCPs. ^9^ The presence of clustered lesions accentuates even more Ap sites reactivity in NCP, with rates 2.5-fold higher than for isolated lesions. ^10^ A particularly deleterious chemistry to be scrutinized arises from the interaction with the numerous positively-charged residues (lysines and arginines) of histone tails. The propensity of Ap sites to form DNA-protein cross-links within NCPs has been thoroughly investigated and quantified by Greenberg and collaborators, ^8,9,11–14^ shedding light on the effect of the superhelical location (SHL) of the abasic site on the kinetics of Ap disappearance and single strand breaks (SSB) formation. ^9^

When located at the mutational hotspot SHL 1.5 of the NCP, isolated and clustered Ap sites react almost exclusively with histone H4, which is responsible for 85-90% of the DPCs in this region. ^8–12^ Even upon deletion of the H4 19 latest amino acids, this histone remains the major player in DPCs formation with Ap89 (80%) and Ap207 (60%). The H3 tail is then involved in 15% and 35% of the DPCs with Ap89 and Ap207, respectively. Mutagenesis studies highlighted the highest implication of H4 Lys16 and Lys20 in cleavage of Ap and its oxidized derivatives at position 89. ^11,12^ His18 and Lys31 might also play a role in DPCs formation of H4 with Ap89 and Ap207, ^8,9^ respectively. However, H3 residues that could react with abasic sites at SHL 1.5 are unknown.

It is documented that clustered abasic sites can induce drastic structural deformations in short B-DNA oligonucleotides which challenge their processing by repair enzymes. ^15,16^ It has also been reported that isolated abasic sites can locally modify the structure and dynamics of B- and N-DNA. ^17,18^ Yet, the impact of clustered abasic sites on a NCP structure remains elusive. Recently, Kurumizaka’s group published the very first X-Ray structures of lesion-containing N-DNA, ^17,19,20^ while the growing number X-Ray structures of canonical N-DNA and its variants allowed to stimulate computational studies. ^21–24^ The response of histone tails in proximity to embedded DNA lesions, either positively-charged or after post-translational modifications, appears important. ^25^ In this work, we investigate histone tails interactions in proximity of cluster abasic sites, which are decisive as they prefigure the formation of DNA-protein crosslinks.

Owing to a simulation protocol we successfully developed and used on isolated, clustered and tandem Ap sites in short oligonucleotides and isolated Ap-containing NCP, ^15,16,18,26^ we report here a mapping of the histone-DNA interactions in a NCP harboring two closely-spaced Ap sites at SHL 1.5 along four micro-seconds molecular dynamics (MD) simulations replicas. ^27^ Our model system, derived from the X-Ray structure of Richmond et al., ^28^ harbors *in silico* mutated Ap sites at positions 89 and 207 (SHL 1.5), which are directly related to experimental data ^8,10^ – see Figure 1. Noteworthy, H4 and H3 tails are truncated in the crystal structure (at positions 15 and 19, respectively), and were kept as such in our simulations. In the starting structure, the H3 and H4 truncated tails are 20-30 Å away from the damaged sites. Four replicas of 1 to 2 *μ*s all-atom classical MD revealed different patterns of non-covalent interactions between the histone tails and both Ap sites, and provided an all-atom description of clustered Ap sites structural behavior in NCPs.

In one MD replica, the H4 tail rapidly folds onto the DNA double-helix in the vicinity of Ap89. Several amino acids surround the Ap site in a very stable hydrogen bonding network – see Figure 2-A and C. Lys20 interacts with the negatively-charged phosphate in the minor groove, while Arg17 intercalates within the major groove, in the gap formed by the absence of nucleobase at the Ap site base-pair. Besides, His18 also places itself very close to the abasic reactive C1’ atom. Arg17, His18 and Lys20 all exhibit amino groups prone to react with the abasic site Ap89. Arg19 and the N-terminal Lys16 are not situated far from Ap89, but interact only transiently with the lesion during the simulation. In the end, Arg19 points towards the solvent and Lys16 interacts with the DNA backbone 3-4 base-pairs below. Interestingly, we observe a similar proximity of the H4 tail in one of the two control MD simulations of the undamaged NCP, although there is no intercalation of Arg17 for there is no gap in the double-helix. The stable interaction between Lys20 and the Ap89 phosphate is also retrieve in this control MD – see Figure S1.

**Figure 2.**
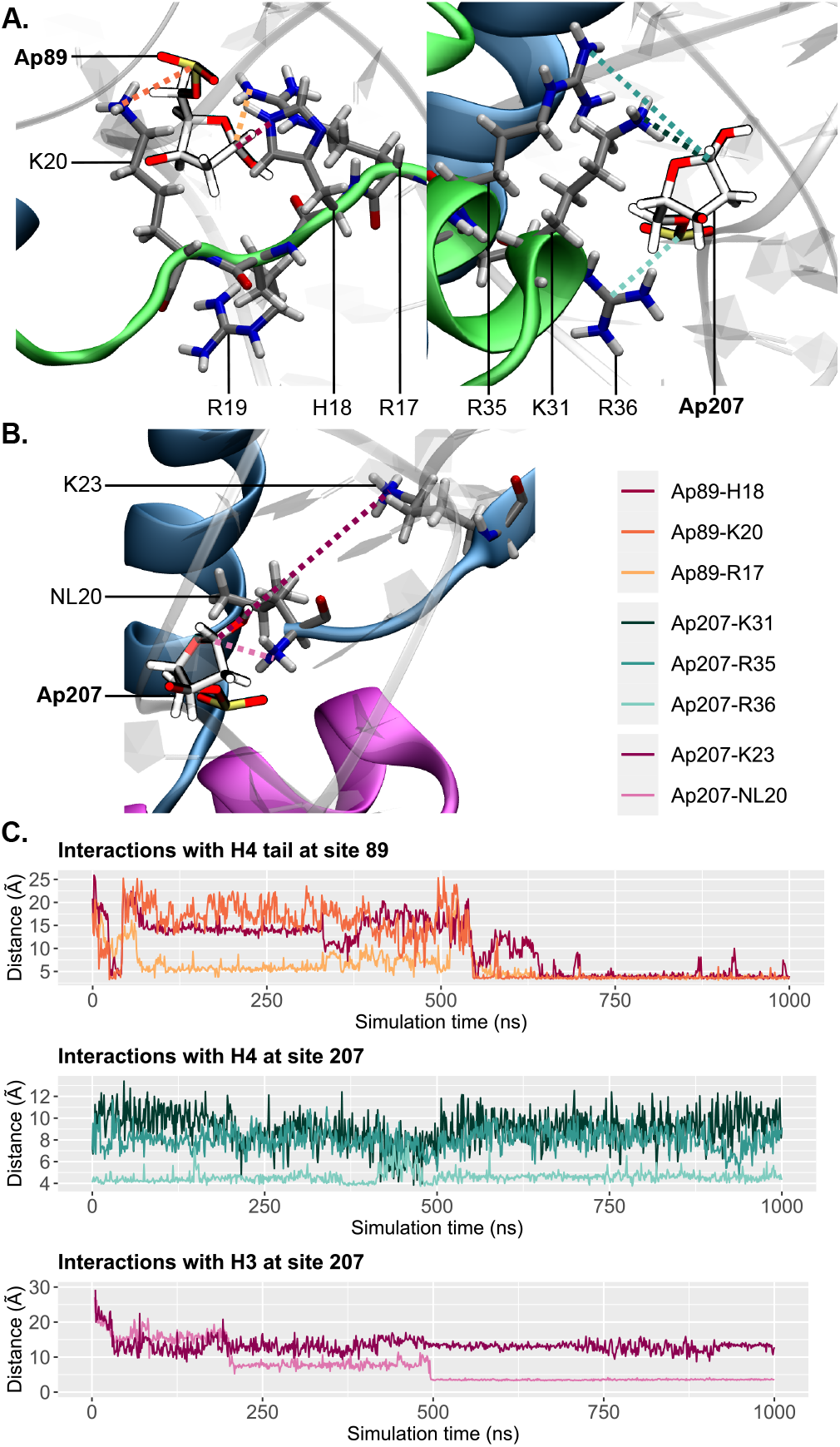
H4 and H3 tails interactions with Ap sites. (**A**) Interactions between the H4 tail and Ap89 involving Arg17, His18 and Lys20 (left), and between Ap207 and the H4 residues K31, R35 and R36 (right). (**B**) Interactions between the H3 tail (N-terminal NL20) and Ap207. (**C**) Key distances between the Ap sites and the histones H4 (top and middle) and H3 (bottom) residues. The color code is as depicted in the legend and in **A**and **B**.

Apart from its tail, H4 forms interactions through its Lys31, Arg35, and Arg36 residues, which are located in the *α*-helix just before the tail. These residues are proximal to the C1’ atom of the Ap207 in all the MD replicas and also in the control simulations. Ap207 can flip towards the histone core during the simulation and interact closely with the H4 core residues – see Figure 2-A and C. Arg36 interacts very strongly with the Ap207 backbone, the typical average distance between its amino groups and the negatively-charged phosphate lying at 4.5±0.5 Å. Its amino groups can also be found very close to the C1’ atom: in MD replica 3, it is found below 6 Å for 25% of the simulation time.

Noteworthy, Sczepanski et al. hypothesized that Lys31 could be involved in DPCs formation for even after removal of its tail, H4 is still responsible for 60% of DPCs formation with Ap207. We show here that the dynamics of the NCPs in the presence of clustered lesions indeed allow a proximity between an abasic site at position 207 and Lys31, but also Arg35 and Arg36 that could be prone to DPCs formation. Such interactions are systematically observed in all the MD replicas with clustered Ap. They are less pronounced in the control undamaged simulations, in which Lys31, Arg35 and Arg36 lie at 10.4±1.2 Å, 12.0±1.5 Å and 8.1±0.2 Å of dA207 C1’, respectively, for the canonical nucleotide cannot flip towards the histone core as the Ap site does.

The H3 tail is responsible for 35% of the DPCs with Ap207 upon removal of the H4 tail. Although the H3 tail is also truncated after Leu20 in our simulations (as it is in the crystal structure), we observe its rapid interactions with Ap207 in one of the replicas. The H3 tail folds near the DNA major groove, in between the two DNA helices turns of the NCP in the vicinity of the lesion sites. The N-terminal NLeu20 rapidly comes close to the C1’ atom of Ap207 (3.5±0.1 Å) with which it forms a very strong hydrogen bond – see Figure 2-B and C. One can surmise that after the removal of the Ap89 site by H4 tail, the remaining SSB might accelerate the reactivity of the proximal Ap207, as already shown in the literature. ^8^ Thus, the H3 tail which is quite close to the Ap207 could come to interact through the major groove and proceed to the second cleavage, leading to the formation of a DSB. This abasic site exhibits a negligible percentage of extrahelicity, although it is partially flipped towards the histone core. Considering the conformational organization of the H3 tail and its folding towards the lesion site, one could postulate that the Lys19 adjacent to the Leu20 (the N-terminal residues in our model) is a good candidate for DPCs formation with Ap sites at position 207. Lys23 appears to stay too far from the lesion site in our simulations – see Figure 2-C. H3 tail harbors other lysines at positions 15, 10, and 5. As the H3 tail is responsible for 11% of DPCs formation with Ap89 even if the H4 tail is present, we can reasonably hypothesize that it might eventually fold over the double helix, rendering the position 89 accessible to Lys5 and Lys10.

The impact of clustered abasic sites on the N-DNA structure has not been investigated so far. Our simulations allow to report the footprint of such deleterious lesions in a NCP. The extrahelicity, a most important structural parameter of abasic sites, ^15,29^ and the bend angle of the 12-bp section harboring the two damaged sites have been assessed to delineate how the NCP structure is impacted by the presence of the clustered lesions. We also quantified the flexibility of the NCP using a machine learning protocol as we previously did for isolated Ap sites in NCP. ^18^

Interactions with the histone core restraint the lesions degrees of freedom, and drastically decrease their extrahelicity compared to naked B-DNA – see Table 1 and distribution in Figure S2. The nucleosomal embedding tends to reduce even more the extrahelical character when the damage faces the histone core (Ap207). Along MD1 and MD2, in which histone tails interact with the lesions, the extrahelicity of Ap89 and Ap207 is of 13.8-18.4% and 0-0.4%, respectively, while in MD3 and MD4 they are of 37.5-38.0% and 5.3-26.3%, respectively. Noteworthy, in MD3 the Ap207 forms stable interactions with H4 Arg36 for 25% of the simulation time, which rationalizes its lower extrahelicity. This suggests that direct interactions with histone tails constrain the Ap sites flexibility by trapping them in stable hydrogen bonds networks. The proximity of the two lesions somehow induces a correlation between their extrahelicities especially when histone tails interact directly within the minor or major groove (see MD1 and MD2). In our previous work on such lesions in B-DNA, we showed that Ap sites adopt an independent structural behavior when separated by more than 4 base pairs, ^15^ which is not the case here.

**Table 1.**
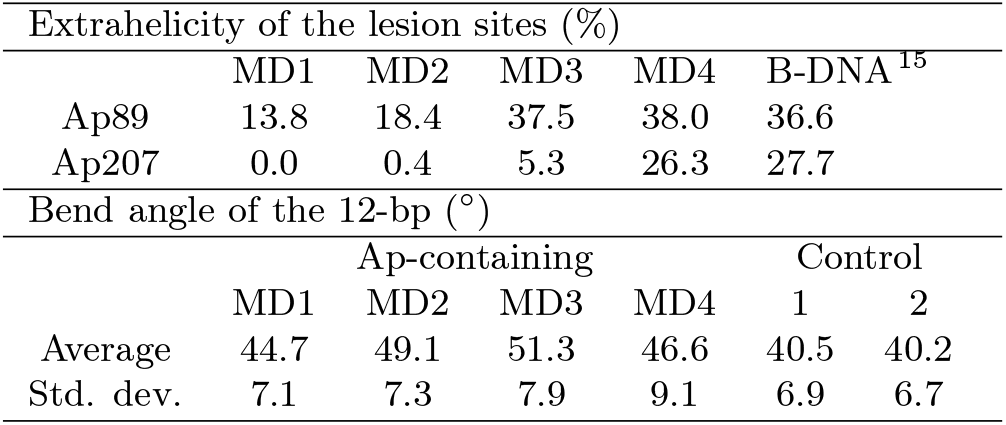
DNA parameters of the 12-bp section harboring Ap89 and Ap207. Extrahelicity values of the Ap sites in N-DNA and B-DNA, ^15^ and averaged total bend angle and standard deviation values for the four MD replicas (MD1-4) and the two control MD of undamaged systems (Control 1 and Control 2).

Unlike in naked B-DNA, no strong perturbation of the adjacent base pairs is induced by the presence of the clustered Ap sites. The nucleosomal embedding maintains the global structure of the double helix and only very local deformations can be pinpointed. Transient re-organizations of the Watson-Crick pairing with base-pairs adjacent to the damage sites can be observed, which palliate the local destabilization resulting from the gap left by the nucleobase removal at Ap sites. Such deformations remain very shy in N-DNA compared to B-DNA, for the nucleosome offers extra-stabilizing interactions. They are mainly focused around Ap207, which tends to flip in and out of the helix, distorting the local helix so that dA85 stacks onto dT208, sometimes leading to the exclusion of dT86 – see Figure S3. Interestingly, the presence of clustered abasic sites accentuates the bend angle of the local 12-bp, with values up to 51.3±7.9° when an undamaged NCP bending lies around 40.5° at SHL 1.5 – see Table 1.

In order to further quantify the changes in flexibility induced by the presence of the clustered Ap sites, we relied on a machine learning protocol that we recently applied for studying isolated abasic sites in N-DNA. ^18^ Probing the contribution of each residue to the overall dynamics of the NCP highlights the increased flexibility of the 12-bp portion of DNA helix that harbors the two lesions – see Figures 3 and S4. Although the abasic sites behavior is more monotonous than in naked B-DNA, the dynamics of the double helix are perturbed by the presence of the clustered lesions. The contribution of the 4 base pairs portion including Ap89 and Ap207 is higher than in the undamaged control system, with contributions up to 10% around Ap207. Ap89 can also show higher dynamics compared to the undamaged system (contribution near 0%), yet less pronounced than Ap207 for it does not induce noticeable structural rearrangements in its vicinity. As one could expect, the highest contributions come from the disordered histone tails, which are highly dynamical along the simulations. This suggests that although the Ap sites can be very locally constrained by the interactions with the histone core and tails, the impact of their presence as clustered lesions induces an increase of the local dynamics of the helix within the NCP. This flexibility might be an important structural signature for lesion recognition and processing by the DNA repair enzymes, which are also promoted by the presence of the characteristic bulges in the DNA backbone near the Ap sites. This supports the possible correlation between local deformation and increased Ap reactivity when clustered, as recently hypothesized by Yang and Greenberg. ^10^

**Figure 3.**
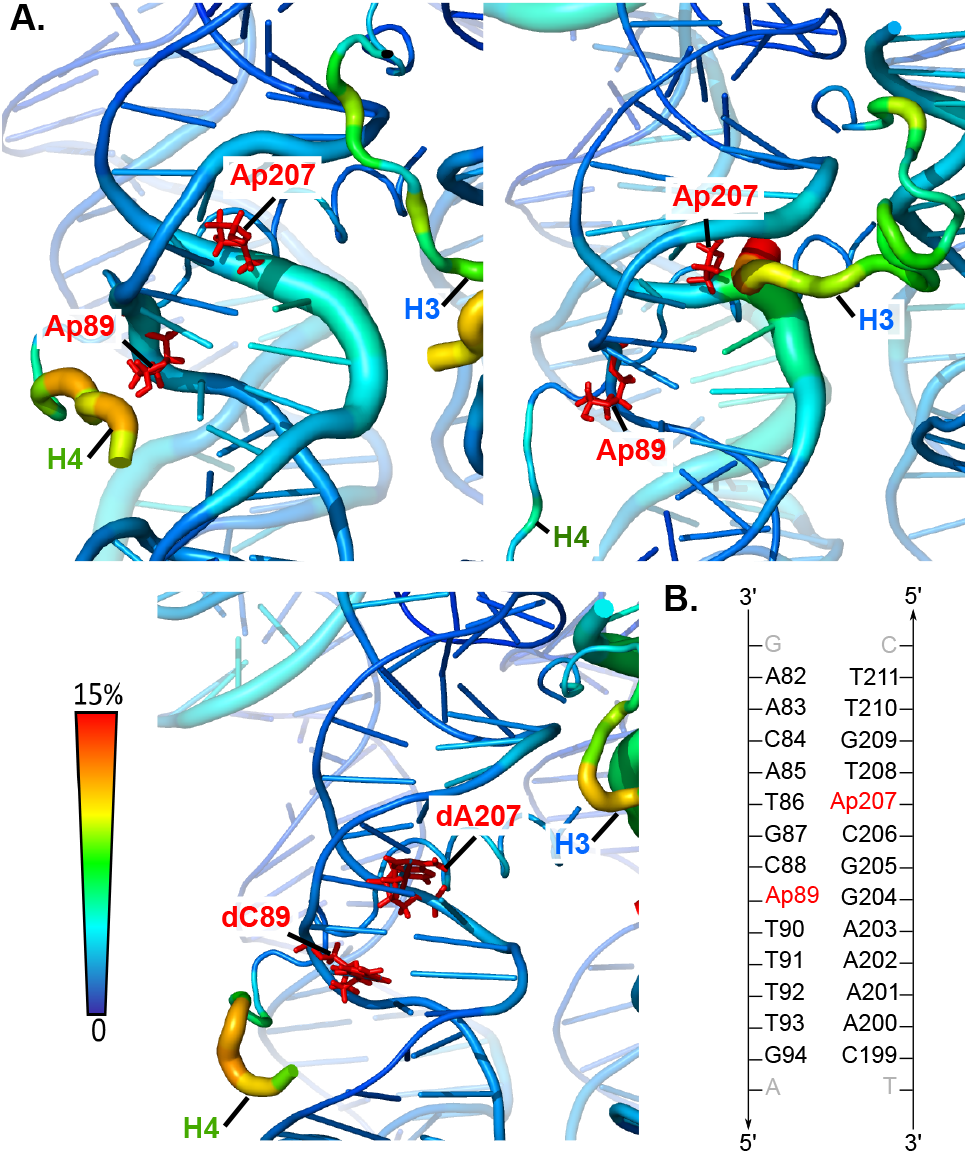
N-DNA structural fluctuations upon presence of clustered Ap at SHL 1.5 based on a machine-learning protocol (see SI). (**A**) Worm representation of the 12-bp harboring the lesions (in red) at SHL 1.5, for simulations replicas in which H4 tail (top left) or H3 tail (top right) interact, and for a control undamaged NCP simulation (bottom). The thickness of the worm and the color scale show the percentage of contribution of the residue to the overall dynamics of the system. (**B**) Sequence of the 12-bp embedding Ap89 and Ap207.

The present study offers unprecedented insights into mapping of complex non-covalent interactions between Ap sites and histone tails. The structural signature of clustered abasic sites within NCP is delineated, and shows contrasted pattern compared to B-DNA embedding due to the interactions with the histone core and tails. ^15,16^ Local reorganization of the double-helix structure is observed upon presence of clustered Ap, although it is much less pronounced than in naked B-DNA. Our unbiased simulations capture the proximity of the H4 tail Lys20, His18, Arg17 and Lys16 to Ap89, and rationalize mutagenesis experiments. Lys20 seems especially crucial as it forms very stable interactions with Ap89 phosphate, which is also observed in simulations of the undamaged NCP. Other residues of H4 (Lys31, Arg35 and Arg36) are found to lie close to the Ap207 and might be involved in DPC formation at this position. Besides, as the N-terminal NLeu20 of the H3 tail can form interactions with Ap207, one can postulate that in the full-length tail, Lys19 is a very good guess for reactivity with Ap sites at this position. The fact that we observe only one histone tail-Ap interaction at a time is not surprising, since the cleavage of one of the damage sites is supposed to induce the formation of a SSB, this latter accelerating the cleavage of the second abasic site. The reactivity of Ap sites upon histone tails removal is decreased, ^8^ yet there are strong interactions already in this scenario, which opens very interesting perspective towards the study of full length histone tails interactions with Ap sites in NCPs. We hone a robust methodology for modeling full length histone tails, which is a timely line of research ^30^ as the community would benefit from extensive benchmarks to identify force fields able to accurately represent such complex combination of ordered and disordered assemblies. The combination of enhanced sampling methods and machine learning algorithms can also be explored to correctly sample the highly complex conformational landscape of the full-length disordered tails, and gives promising openings towards further studies. This work constitutes a proof of concept for studying DNA-protein interactions within a nucleosome core particle harboring multiple sites damage, and our protocol sets the grounds for further studies of Ap reactivity upon histone post-translational modifications ^31^ and the role of histone tail in DNA repair proteins recruitment. ^32^

## Acknowledgement

The authors are grateful to ENS de Lyon for support and the Pôle Scientifique de Modélisation Numérique for calculation resources. This work was performed within the framework of the LABEX PRIMES (ANR-11-LABX-0063) of Université de Lyon, within the program “Investissements d’Avenir” (ANR-11-IDEX-0007) operated by the French National Research Agency (ANR).

## Supporting Information Available

Details of the computational procedure; supplementary figures; movies of the MD trajectories exhibiting the rapid interaction of H3 and H4 histone tails with the lesion sites. These files are available free of charge.

---

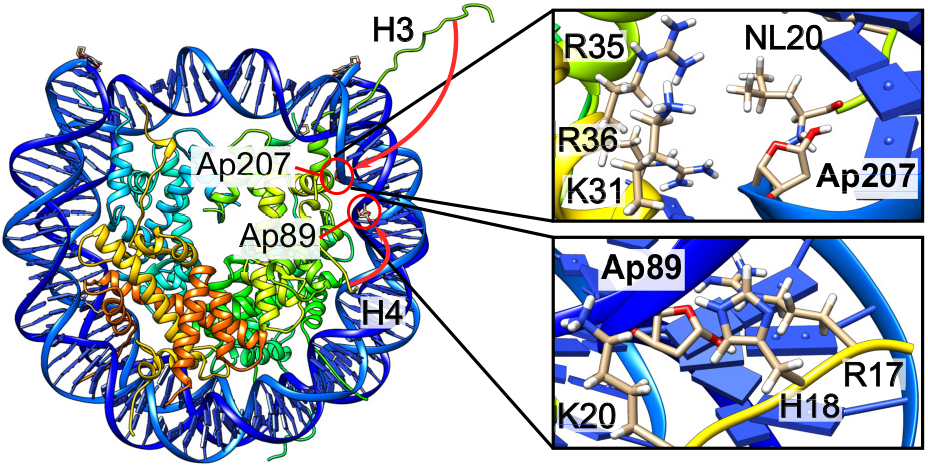
For Table of Contents Only

## Notes

### Competing Interest Statement

The authors have declared no competing interest.

