## Supporting Information for "A dynamic view of histone tails interaction with clustered abasic sites in a nucleosome core particle"

### Computational details

**System setup** All classical MD simulations were performed with the Amber18 suits of programs,[1] using the parm14SB force field[2] with the bsc1 corrections for DNA backbone.[3] Abasic site parameters were generated as described in our previous study.[4] Abasic sites were added *in silico* on positions 89 and 207 of the $\alpha$-satellite 146-bp N-DNA structure reported by Richmond and coworkers[5] (PDB ID 1AOI). The crystallographic structure exhibits portions of the histone tails: 3, 23, 19, and 15 amino acids missing for H2A, H2B, H3, H4 respectively. Because simulating the full length disordered tails would cause problems in terms of force field accuracy and time sampling in our calculations, tails were kept truncated. Amino acids were protonated using the H++ server (3.2 version - http://biophysics.cs.vt.edu/H++),[6–8] and the NCP was soaked within a truncated octahedron-shaped TIP3P water box with a 12 Å buffer. Potassium cations were added to ensure neutrality, resulting in a system of $\tilde{1}95{,}000$ atoms.

**Molecular Dynamics simulations** The starting structure was first optimized in a 10,000 steps run. Temperature was then increased from 0 to 300K and kept stable for the remaining of the simulation using Langevin thermostat with a $\gamma$ln collision frequency of 1 ps$^{-1}$. The system was then equilibrated for 2 ns prior to the 1-2 $\mu$s production run. Four replicas were performed in order to enhance the sampling efficiency. The same protocol was carried out on control undamaged NCP systems, with two 1$\mu$s MD replicas.

**Structural analysis** Structural analysis was performed using Curves+[9] on the 12-bp portion harboring the clustered lesions. Extra-helical character of the AP sites was monitored as described by Lavery and coworkers.[10] The clustering and distance analysis was performed using the cpptraj module of AmberTools18.[1] The contribution of each residue to the overall NCP dynamics was quantified using a machine learning protocol based on a Principal Component Analysis (PCA) using the internal coordinates (inverse distance between geometric centers of two residues) for protein and DNA residues along the trajectory as input to build a covariance matrix, as already done for single Ap damages.[11] The PCA was performed with a home brew script utilizing the Scikit-learn58library.[12] To obtain the per residue importance, the sum of the weighted principal components up to certain threshold with the corresponding as weights is taken and then normalized.

### Figures

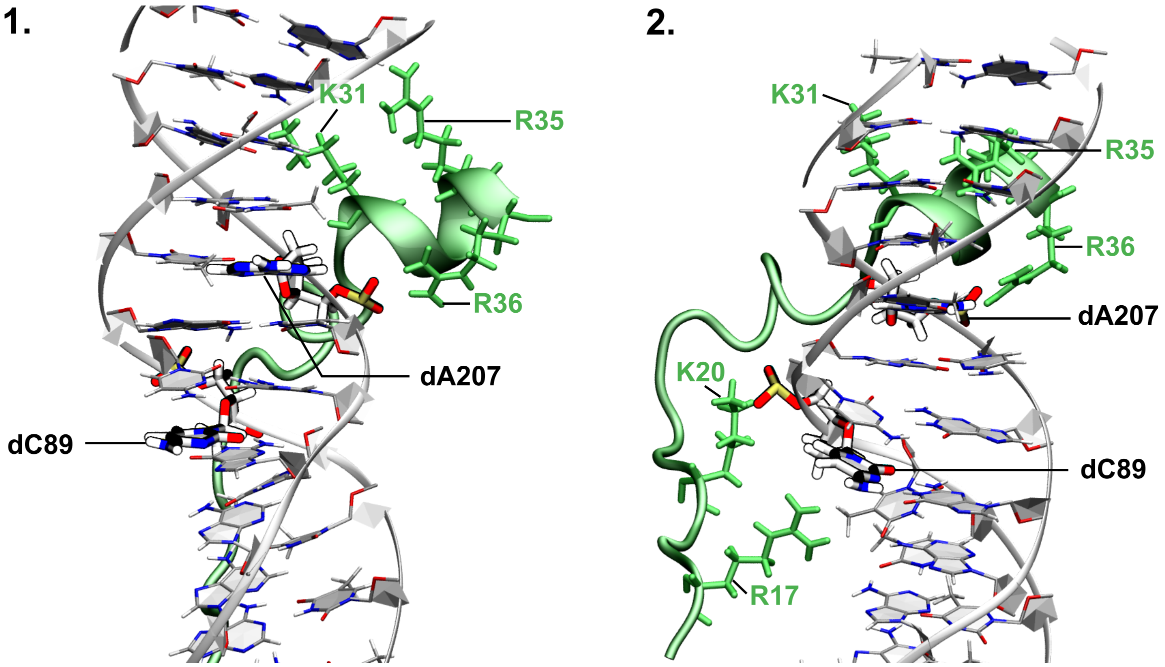


Figure S1 – Representative structure of the 12-bp in organization around position 89 and 207 in the two MD replicas (1 and 2) of the control undamaged system. The histone H4 tail appears in green.

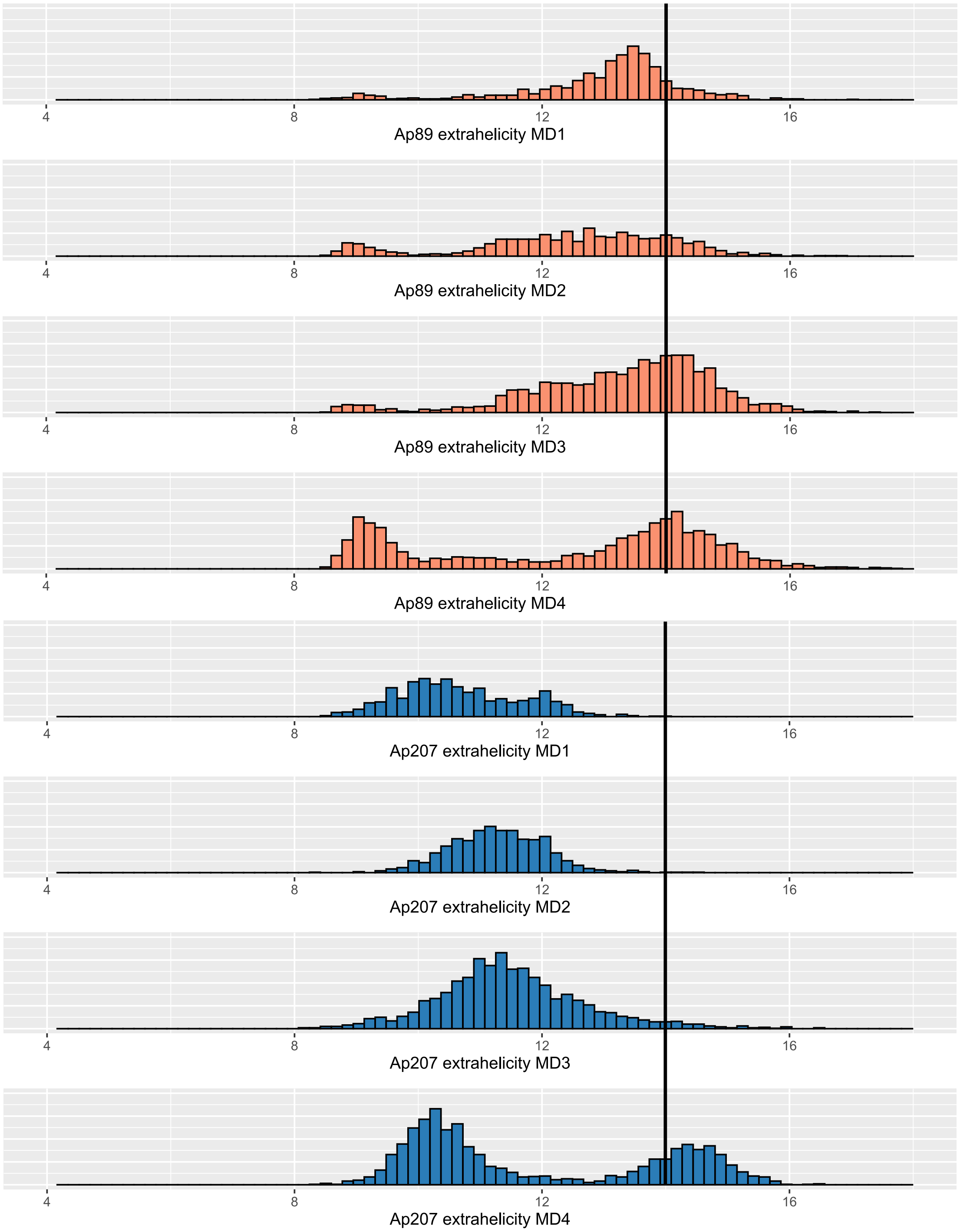


Figure S2 – Distribution of the distance between the Ap site C1' atom and its facing orphan base C1', at position 89 (orange) and position 207 (blue) over the four MD replicas. Above the threshold of 4Å, the abasic site is considered as extrahelical.

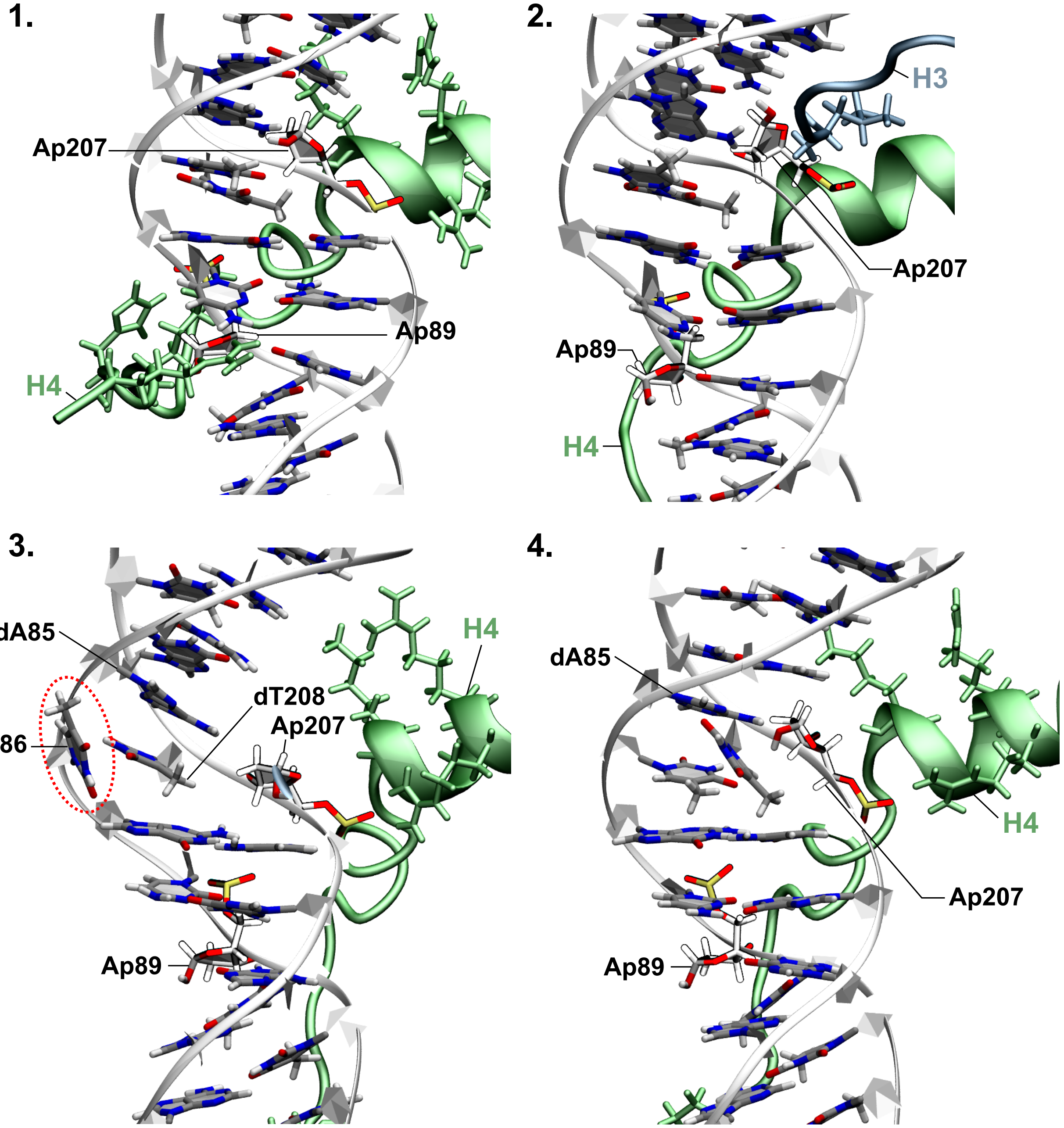


Figure S3 – Representative structure of the 12-bp in organization around the two lesion sites in the four MD replicas (1, 2, 3, 4). The histone tails appear in green (H4) and blue (H3).

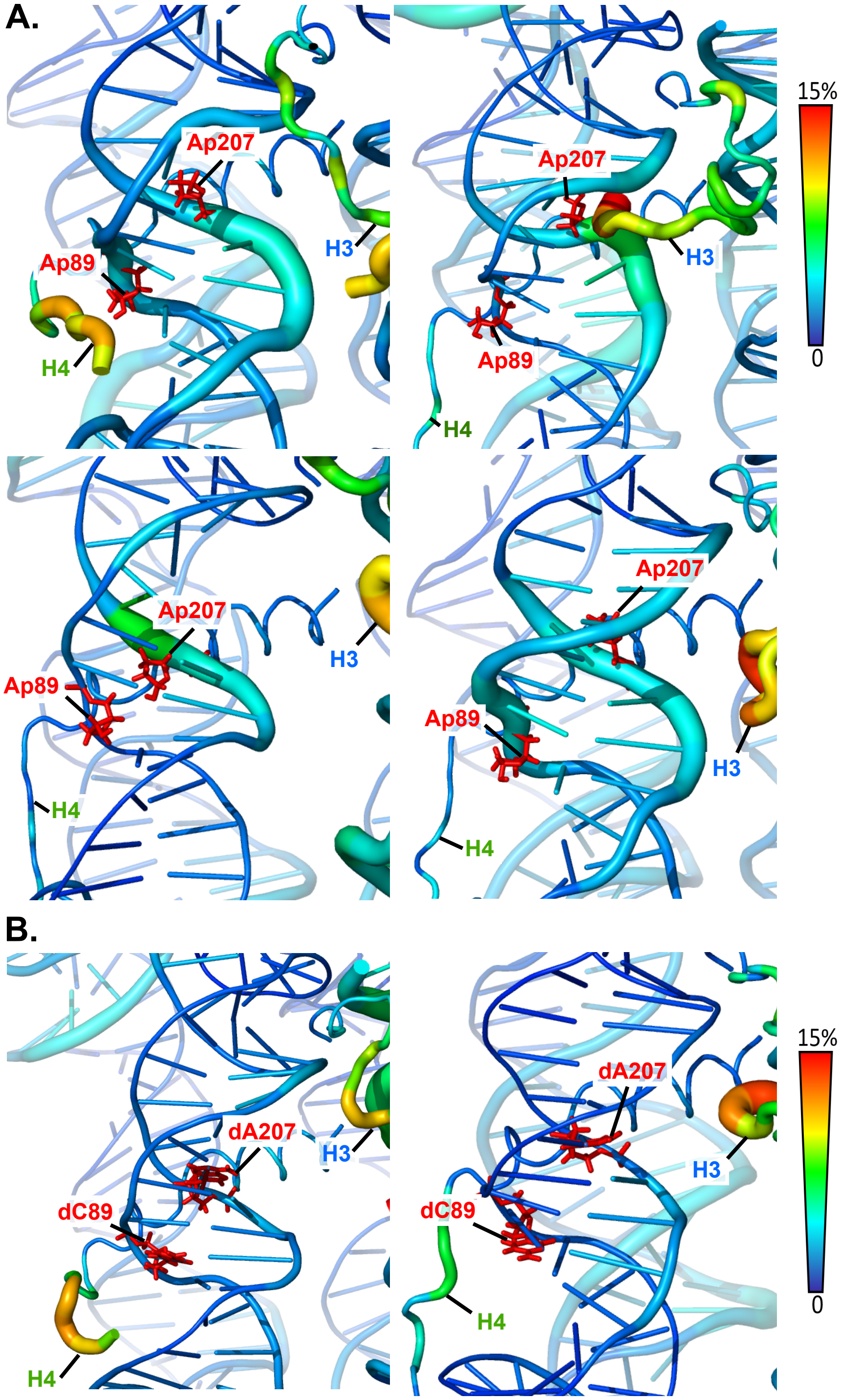


Figure S4 – PCA of N-DNA structural fluctuations upon presence of clustered Ap at SHL1.5. (**A**) Worm representation of the 12-bp harboring the lesions (in red) at SHL1.5, for simulations replicas 1 (top left), 2 (top right), 3 (bottom left), 4 (bottom right) and (**B**) replicas 1 (left) and 2 (right) of the control undamaged system. The thickness of the worm and the color scale show the contribution of the residue to the overall fluctuations of the system.
